# Feature selection for the classification of fall-risk in older subjects: a combinational approach using static force-plate measures

**DOI:** 10.1101/807818

**Authors:** D’Ó’ Reilly

## Abstract

**Introduction:** Feature selection prevents over-fitting in predictive models. This study aimed to present an effective feature selection method that leads to a reliable classification of fall-risk in older subjects using static force-platform data across four conditions only.

**Method:** 528 features were generated from a publicly available dataset of force-plate signals from 45 low-risk and 28 high-risk subjects. Subjects were classified as high- or low-risk if they recorded ≥1 falls in the prior 12 months and/or were rated as high-risk on the FES. The feature selection protocol included SVM-RFE, GA and ReliefF and finally SAFE. Several machine-learning models were then used to evaluate classification performance.

**Results:** 67 features were identified after the three-fold process which was further reduced to 18 features after SAFE. The MLP achieved the highest average classification accuracy of 80%. All classification models evaluating this final subset displayed high variance across all performance metrics, especially in terms of sensitivity to high-risk subjects.

**Interpretation:** An optimal feature set of static force-plate measures was insufficient in creating a reliable classifier of fall-risk. This was due potentially to the limited information about fall-risk that could be provided by such measures leading to under-fitting/over-fitting being unavoidable and appeared to be centered around an insensitivity to high-risk subjects.

**Conclusion:** Static stability measures have shown some usability in fall-risk classification however feature sets limited to such measures are inadequately sensitive to high-risk subjects. The utilized feature selection methods demonstrated their ability to identify relevant stability measures and could be used successfully on dynamic measures.

## Introduction

Feature selection as a method for identifying optimal variable subsets has come into the spotlight in the age of ‘*big data*’. This is due to the numerous advantages provided by selecting an optimal set of features rather than using huge datasets including higher accuracies, lower computational loads and the prevention of over-fitting (1). Feature selection methods can be categorized into filter, wrapper and embedded methods. Filter methods incorporate ranking techniques that order features in terms of discriminative power, wrapper methods rate features based on their predictive performance and embedded methods make use of classification models and can be carried out simultaneously during the training process (1–3). Traditionally an optimal set of features was said to be one that contained the most relevant features available (i.e. provided the most information about the target) (2). What is considered an optimal feature set has evolved from this and thus the coinciding measures of feature suitability have also. Yu (4) discusses in detail how relevancy of features can be broken down into levels (Strongly relevant, Weakly relevant but non-redundant, Weakly relevant and redundant and irrelevant). This perspective also states that relevancy analysis alone can be regarded as inadequate without redundancy analysis also. Two features are considered completely redundant to each other when one does not contribute any unique information to a target variable when compared to the other (4). The presence of feature redundancy should be minimised as losses in information can occur leading to sub-optimal model performance. Redundancy occurs when features are highly correlated but can still have an effect at lower correlations. Having all features completely orthogonal is an unlikely scenario in most cases and therefore some redundancy is unavoidable. Complementarity is a third element in feature selection that focuses on the information gained via synergies between features (5,6). Some features taken alone may not be highly relevant but together with other features can provide substantially greater information overall..

In motion analysis research, as is the case in many biomedical fields, a significant number of features can be gathered from a single individual while samples are often difficult to come by (7). This creates a situation where the number of features is far greater than the number of sample participants, an unfavourable situation when constructing models as over-fitting is likely. A current societal issue being tackled by motion analysis research is that of falls in at risk populations such as the elderly and neurologically impaired (8–10). In the healthy older population, falls can occur in individuals with no obvious balance impairments and can be relatively independent of age in this population (11). This makes the early identification of at-risk individuals a crucial avenue of research. Stabilometry is the study of the postural control system and assessments take place during dynamic movements such as gait or in a static, quiet stance position (12,13). Functional balance measures such as the “Berg balance scale” and “Timed-Up & Go” test possess a ceiling-effects when assessing active older subjects and are not sensitive to early, subtle deteriorations in balance mechanisms (14,15).

The use of a force-plate has become common place in stabilometry as highly informative and reliable signals can be generated (16). Measures of COP derived from a force-plate for example are capable of differentiating subjects in terms of age-group and between fallers & non-fallers (17,18). Stimulating the postural control system by creating challenging conditions while on a force-platform such as with compliant surfaces, narrowing-stance, closing eyes etc. have also been effective in differentiating fallers from non-fallers (17,19,20).An extensive range of static force-plate measures have been found to be relevant to fall-risk assessment including COP velocities, RMS of COP displacement across various conditions among others (21). However no one measure can be deemed a blanket predictor of fall-risk (21,22). Many of these studies also have conflicting evidence for the predictability of these measures in terms of fall-risk (21). There are a number of potential reasons for this conflict including the heterogenous nature of and causes for balance deficits in old age and disorders common in older populations creating additional variability in samples (23,24). It is potentially the case that an optimally selected set of these measures may classify individuals in this population with a more reliable accuracy.

Recent studies on fall-risk predictors have recognised the necessity for multiple features and non-linear algorithms in classifying fall-risk by exploring the potential of machine-learning to create effective classification models (25–29). The majority of these studies have focused on dynamic movement analysis to predict elderly fall-risk (25,26), while the presence of a thorough feature selection process has been lacking with one study of note using principal component analysis (30). The aim of this study is to demonstrate an effective feature selection procedure that can allow for more accurate and reliable classification of fall-risk among older subjects using static force-platform measures only. This study will avail of an opensource dataset (https://peerj.com/articles/2648) which has been published specifically for use in such studies (31). A combination of Support vector machines-Recursive feature elimination (SVM-RFE), a Genetic algorithm (GA) and ReliefF feature selection methods will be utilized to find the most relevant, least redundant subset and Singha & Shenoy’s (6) self-evaluating feature evaluation (SAFE) heuristic will optimize complementarity downstream from this. A combinational approach to feature selection has demonstrated notable advantages over single feature selection methods (4,32). The performance of the final feature subset will then be tested using a Multi-layer perceptron (MLP), Support vector machines (SVM), Naïve Bayes (NB) and K-Nearest neighbours (K-NN).

## Methods

### Force-platform dataset

Force-platform data including COP coordinates, GRFs and Torsional forces (Moments) in all directions was acquired via an open source repository published by Santos & Duarte (31) who analysed the postural stablity of older subjects during quiet stance throughout four conditions (Eyes open/closed on firm surface & Eyes open/closed on a compliant surface). A full description of the data acquisition protocol can be found at (31). Only the most relevant aspects to this study will be covered here. Subjects met inclusion for this study if they did not have a serious physical disability (e.g. neurological impairment). To maintain representativeness, subjects met inclusion if they had common musculoskeletal disorders such as Osteoporosis for example. Table 1 below illustrates the characteristics of this sample group.

**Table 1:** *Significant features (p<0.05)

| *Fall-risk Groups (n=73)* | *Low Fall-risk* | *High Fall-risk* | *Mann Whitney-U test* |
| --- | --- | --- | --- |
| **Gender** | M=12/F=33 | M=2/F=26 | - |
|  | **Mean (SD)** | **Mean (SD)** |  |
| **Age (years)** | 72 (6.5) | 71(6.3) | ($p=0.5$) |
| **Height (m)** | 1.58 (9.1) | 1.55 (5.9) | ($p=0.5$) |
| **Weight (kg)** | 64 (8.3) | 62 (8.5) | ($p=0.5$) |
| **BMI (kg /m<sup>2</sup>)** | 25.5 (2.9) | 26 (2.9) | ($p>0.5$) |
| **Foot length (cm)** | 23 (1.3) | 22 (1.3) | ($p<0.05$)* |
| **No. of medications** | 2.4 (1.5) | 2.3 (1.8) | ($p>0.5$) |
| **No. of disorders** | 1.02 (0.97) | 1.24 (1.1) | ($p>0.4$) |
| **Falls (Past 12 months)** | 0 (0) | 2.5 (9.4) | - |
| **FES (Short version)** | 9.2 (1.7) | 12 (4.2) | ($p<0.01$)* |
| **TMT time A** | 62.5 (54.2) | 53 (22) | ($p>0.5$) |
| **TMT time B** | 167.2 (95) | 174.1 (84) | ($p>0.5$) |
| **BesTest** | 18 (5) | 19 (3.4) | ($p>0.4$) |

Falls history in the past 12-months and risk-rating in terms of the FES were used to categorise the fall-risk groups in this study. If a subject had at least one fall recorded and/or a high-risk rating as per the FES at the time of analysis, they were considered high-risk with the remaining subjects allocated to the low-risk group. 45 low-risk subjects (Age:72±6.5, FES score: 9.2±1.7) and 28 high-risk subjects (Age: 71±6.3, Falls history: 2.5±9.4, FES score: 12±4.2) will be analysed in this study. No significant differences were found between groups in terms of the Trail Making test (TMT A & B) or the BesTest. Significant differences between the groups were present for FES scores (p<0.01) and foot-lengths (p<0.05) only.

### Data acquisition and processing

Subjects balance was analysed barefoot for 60 seconds per trial with three trials recorded per condition. Feet were placed at 20 degrees with the heels separated by 10cm in each trial. This stance was standardized across trials by marked lines on the force-platform (See figure 1). The order of the four different conditions was randomized to offset the effect of practice. Conditions included a combination of visual & surface manipulation. For ‘*Eyes-open*’ conditions, subjects were asked to focus on a 5cm round black target at eye-level on a wall 3 metres ahead of them. During ‘*Eyes-closed*’ conditions, subjects were asked to look at this target initially and then close their eyes once a comfortable position was found. A rigid surface was utilised for the ‘*Firm surface*’ condition while the ‘*Compliant surface*’ condition was conducted using a 6cm high foam block (Balance Pad; Airex AG, Sins, Switzerland).

**Figure 1:**
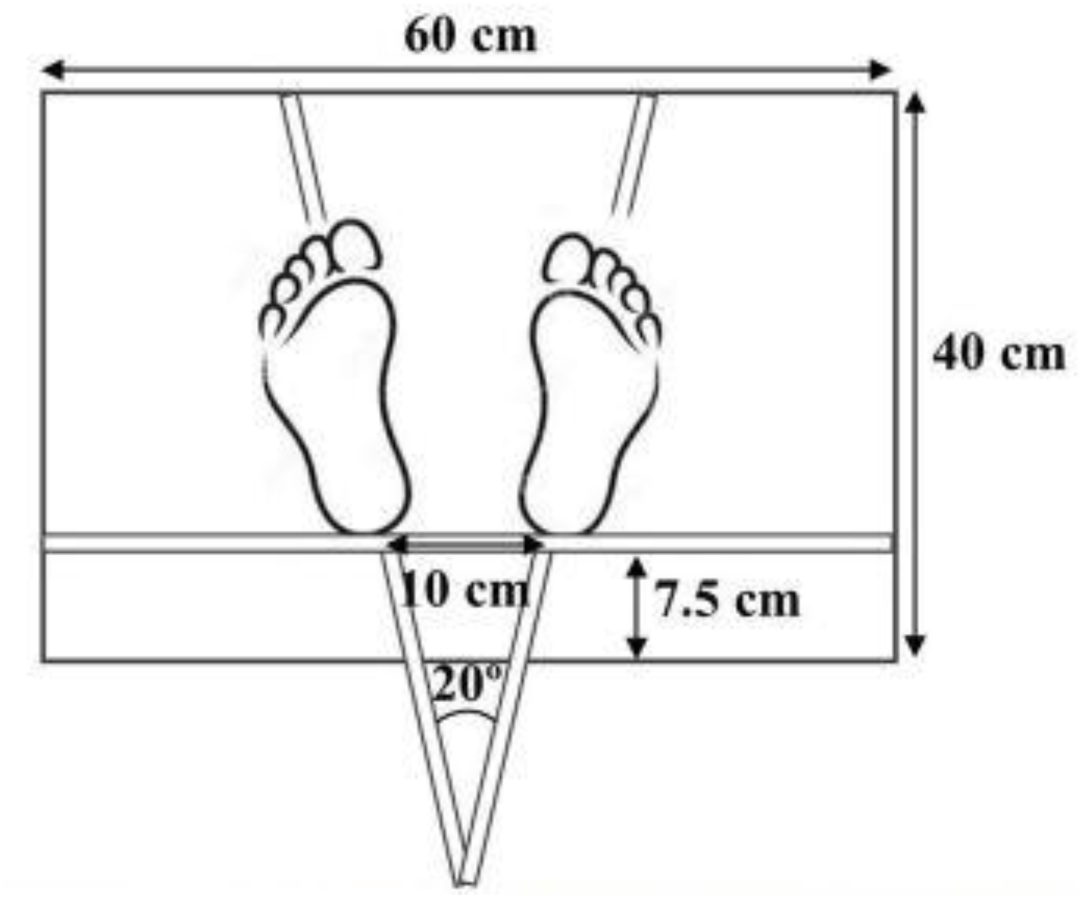
Santos & Duarte (31)

Signals were collected via a 40cmx60cm commercial force platform (OPT400600-1000; AMTI, Watertown, MA, USA) and amplifier (Optima Signal Conditioner; AMTI, Watertown, MA, USA) at a sampling frequency of 100 Hz. The mean COP accuracy of this force-plate was approximately 0.02cm. Data acquisition was performed using NetForce software (Version 3.5.3; AMTI, Watertown, MA, USA). All data was smoothed with a 10Hz 4^th^ order zero lag low-pass Butterworth filter.

### Feature generation

For the purpose of this study, an extensive range of time- and frequency-domain measures were derived from the dataset and summary statistics of these measures including Mean, SD, minimum, maximum, range and RMS were taken for analysis also. Many of these measures can be found in more detail in Prieto et al (13). The same measures from different conditions were considered as a unique feature. This process culminated into a total of 528 features altogether, creating a necessary situation where the sample size n<<p the number of features.

#### Time-domain measures

The COP position, displacement and velocity in AP and ML directions was obtained. All GRFs were normalized by subject weight (kg) while the vertical torque or Free moment (T_Z_) was normalized by weight and height squared as seen in (33). This measure is sensitive to asymmetries in postural control about the vertical axis. The variance signal of T_Z_ can capture specific types of postural adjustment (i.e. slow-leaning motions versus fast corrective changes in posture) when low- and high-pass filtered respectively (36,37). A 5^th^-order digital finite duration impulse response (FIR) filter with a cut-off frequency of 0.1 Hz for both the low-pass (<0.10Hz) and high-pass (>0.1Hz) signals was applied in this case. Phase-plane plots combining COP displacements and velocities vectors were computed in both the AP (*APr*) and ML (*LATr*) directions with a global phase-plane (R) also computed from the resulting variables (35). The COP Path length (PL), Resultant distance (RD), Mean distance (MDIST), RMS distance (*RDIST*), Total excursion (*TOTEX*), mean velocity (*MVELO*), 95% confidence-circle area (*AREA-CC*), mean rotational frequency of the COP (*MFREQ*), Fractal dimension (*FD*) and 95% confidence-circle of the Fractal dimension (*FD-CC*) were also calculated (13).

#### Frequency-domain measures

Frequency-domain features were generated via a customized python script (36). Total power of the frequency spectrum (*P*_total_), mean frequency (*MFreq*), median frequency (*Freq*_50%_), 95% Power frequency (*Freq*_95%_) and Peak frequency (*Freq*_peak_) were extracted (13). Frequency spectrums were derived from both COP displacement and shearing forces in both anterior-posterior and medio-lateral directions. Prior to transforming the COP displacement signal, the COP position was detrended so sub-trends could be properly analyzed. Along with the entire frequency band of the displacement and shearing force signals being investigated, the signals were also filtered to specific bandwidths sensitive to certain postural control systems using a 1^st^-order Butterworth band-pass filter. The frequency band 0.01-0.1Hz is sensitive to the visual control system while 0.1-0.5Hz and 0.5-1Hz are sensitive to vestibular and somatosensory control systems respectively (37).

### Feature selection

As mentioned, a combinational approach to feature selection will be carried out. SVM-RFE is a well-known wrapper feature method that has found wide use in various fields (38,39). Backward feature elimination has been noted as an effective way of removing irrelevant and redundant measures and can achieve good generalization (1,39). Starting with the entire feature set, SVM-RFE will take out one feature at a time that is ranked lowest while the accuracy of the remaining feature set will be 10-fold cross validated. This method will begin the feature selection process by reducing the original feature set into a smaller, more manageable subset of features and reduce the effect of redundant and irrelevant features further downstream in the Genetic algorithm. Genetic algorithms come from a family of algorithms known as ‘Evolutionary algorithms’ that are inspired by processes in nature and generally incorporate the idea of a population of potential optimization solutions with each individual with the population corresponding to a feature subset for example (40). A fitness score is then calculated on the individuals in the population and the next generation of individuals adapts and higher fitness scores are propagated until a specified number of generations have been run and the fittest individual is selected. A genetic algorithm based on the Distributed Evolutionary algorithms in Python (DEAP) framework will be embedded in an SVM classification model (41). The feature subset to come out of SVM-RFE will be input into this embedded feature selection method. A combination of SVM-RFE and GA has successfully been used previously on highly complex biomedical data (38).

Parallel to the above-mentioned process, a filter-based feature selection algorithm called ReliefF will be applied to the original feature set also. This filter-based method is classifier-independent but uses K-Nearest neighbors distance metrics to determine the most discriminative features (3). This method is sensitive to interactions and local dependencies between features and the discriminative power of the feature as a result of this.

Finally the features identified by GA and ReliefF selection will be combined and an optimal subset will be obtained via SAFE, a complementarity-based feature selection procedure proposed by (6). This procedure allows for trade-offs to occur between information lost through redundancy (Rs), information gained through relevancy (As), Dependency between features (Ds) and information gain via complementarity (Cs) to be adaptively assessed as the feature set is iteratively searched through. For further details on the formulation of As, Rs, Cs and Ds please see (6). Equation (1) and (2) below illustrate the SAFE procedure where |S| is the subset size, β is derived from the hyperparameter α that controls the trade-off between redundancy and complementarity while γ is derived from γ which controls the complementarity-relevancy trade-off in the feature subset allowing smaller subsets to be favored. This method does not suffer from initial feature selection bias as each feature is assessed continuously throughout the process meaning the feature subset is less likely to be skewed towards a dominant sub-group in the sample. The number of features to keep in the final subset is often arbitrary or based on expert decisions in feature selection. This is not necessary in the case of SAFE as the search continues until an optimal subset score (S) is reached.

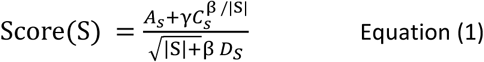

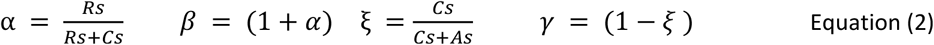

As entropy estimations are necessary for the SAFE procedure, the feature subset to be assessed in this case will be discretized using the Class attribute interdependency maximization (CAIM) method (42). This discretization method was chosen due its ability to preserve target attribute interdependency while minimizing the number of intervals required to describe the data. After the final SAFE procedure has been carried out, the feature subset with the highest (S) score will input in its discretized form into an MLP, NB, K-NN and SVM classifiers to assess performance with the mean, SD and range of score values from 25 rounds presented. These classifiers were chosen based on the contrasting ways they learn the training data. This evaluation and feature selection process described will be carried out using Python and Python’s inbuilt Sci-Kit learn module (43). Classification accuracies, area under the ROC-curve (ROC-AUC), F1 score, precision and recall for the specific risk classes will be presented to illustrate the performance of the classifiers. The code for the SVM-RFE - Genetic algorithm - ReliefF feature selection process described above including all parameter used can be found at the following Github repository: **https://github.com/Davidoreilly12/Feature-selection-study.git**.

## Results

### Combinational feature selection

SVM-RFE concluded with 87 features from the original set of 528 identified as useful for the classification of fall-risk in this elderly cohort. Figure 2 shows how variable the accuracy of a model can behave due to irrelevant/redundant features. The feature set was further reduced to 67 features by ReliefF and the Genetic algorithm. ReliefF identified features from the ‘*Eyes-Open/Firm-surface*’ condition only, many of which are sensitive to COP migration. FD-CC and P_Total**[0.01-0.1Hz]**_**-**COP_X_ were the only features selected in the SAFE feature set to come from ReliefF with the remainder being identified by the Genetic algorithm. The GA was able to identify the fittest individual within 10 generations and achieve a fitness score of 100% with this individual.

**Figure 2:**
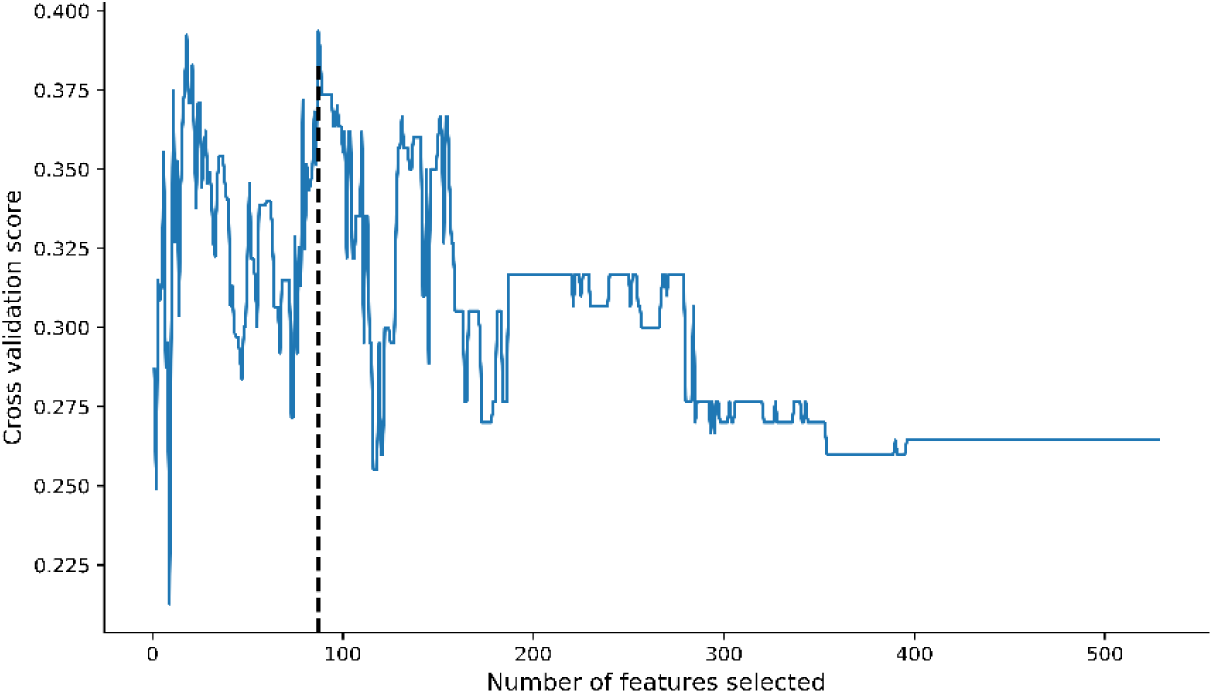
SVM-RFE output (Vertical black line denotes optimal feature subset score)

From the subset of 67 features remaining, a search strategy where the feature with the lowest individual (S) score was iteratively removed until the highest subset (S) score was found. Figure 3 demonstrates the value of feature subset relevancy (As), redundancy (Rs), complementarity (Cs) and dependency (Ds) and how they changed over the course of this process represented in information bits.

**Figure 3:**
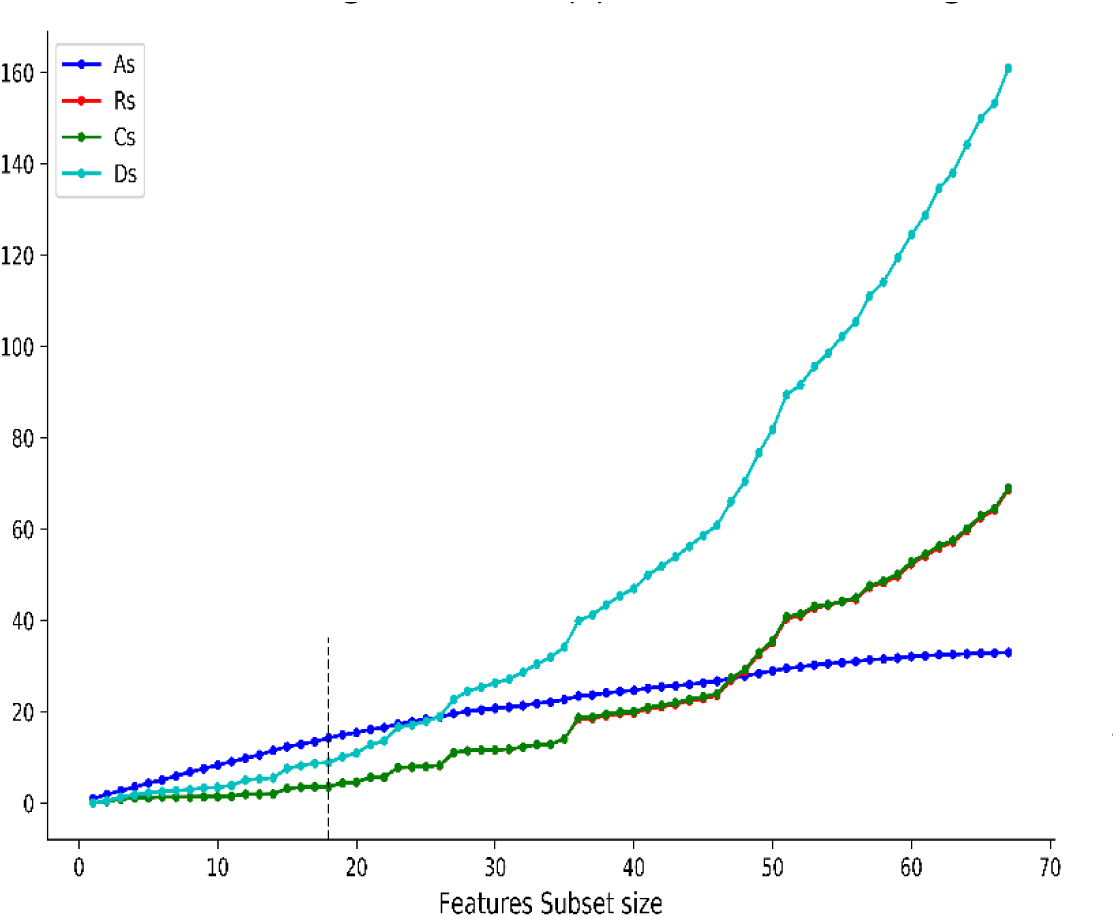
SAFE (Vertical black line indicates optimal feature subset)

An optimal (S) score of 2.705 was found with 18 features. A high feature dependency was present at the commencement of this process and experienced the largest reduction from 161 bits to 8.83 bits in the final subset. Feature subset relevancy (As) was reduced conservatively during the search from 33 bits to 14.2 bits also. The subset was found to have an even amount of redundancy and complementarity present with both Rs and Cs values equal to -/+ 3.55 bits respectively. This was the case throughout the search, effectively cancelling out the information lost by redundancy with the information gained through interactions.

Table 2 conveys the 18 features, their individual (S) score and the p-value derived from a Mann Whitney-U test for significant differences between the high- and low-risk groups for each feature. Only five of the eighteen features in the final subset were found to have a significant difference between the two groups including Maximum and Minimum F_Y_, Freq95%-F_X_, Mean F_Z_ and Freq95%-F_Y_ all of which were sourced from the ‘Closed-eyes/Firm surface’ condition indicating the majority of features were non-linearly related to fall-risk in this sample. The individual (S) scores ranged from 0.19 for Maximum T_Z_ to 0.403 for Maximum F_X_. The higher the (S) score the more complimentary the feature can be considered towards other features in the subset. Features in this optimal subset with low (S) scores could be said to of a high relevancy for fall-risk classification in this sample but did not provide a significant amount of supplementary information through synergies with other features.

**Table 2:** * features with a significant difference between risk-groups

| **Feature** | **Condition** | **(S)** | **Mann Whitney-U test**<br>**For significant differences between groups** |
| --- | --- | --- | --- |
| **Maximum $T_z$** | 'Open-Eyes/Firm surface' | 0.19 | $p > 0.5$ |
| **$F_z$ (SD)** | 'Open-Eyes/Compliant surface' | 0.293 | $p > 0.5$ |
| **R** | 'Open-Eyes/Firm surface' | 0.314 | $p > 0.5$ |
| **$F_z$ (SD)** | 'Open-Eyes/Firm surface' | 0.343 | $p > 0.05$ |
| **$P_{Total}[0.01-0.1Hz]-COP_x$** | 'Open-Eyes/Firm surface' | 0.302 | $p > 0.5$ |
| **Maximum $F_y$** | 'Closed-Eyes/Firm surface' | 0.321 | $p < 0.05^*$ |
| **Minimum $F_y$** | 'Closed-Eyes/Firm surface' | 0.32 | $p < 0.005^*$ |
| **$MFreq_{[0.01-0.1Hz]}-F_x$** | 'Closed-Eyes/Compliant surface' | 0.333 | $p > 0.1$ |
| **$Freq_{Peak}[0.01-0.1Hz]-COP_x$** | 'Open-Eyes/Compliant surface' | 0.36 | $p > 0.1$ |
| **$Freq_{95\%}-F_x$** | 'Closed-Eyes/Firm surface' | 0.364 | $p < 0.05^*$ |
| **FD-CC** | 'Open-Eyes/Firm surface' | 0.33 | $p > 0.5$ |
| **Mean $F_z$** | 'Closed-Eyes/Firm surface' | 0.351 | $p < 0.01^*$ |
| **$Freq_{Peak}[0.5-1Hz]-F_y$** | 'Open-Eyes/Compliant surface' | 0.3 | $p > 0.4$ |
| **$Freq_{95\%}-COP_y$** | 'Closed-eyes/Firm surface' | 0.34 | $p > 0.1$ |
| **$Freq_{95\%}-F_y$** | 'Closed-Eyes/Firm surface' | 0.34 | $p < 0.05^*$ |
| **Minimum RD** | 'Closed-Eyes/Firm surface' | 0.37 | $p > 0.05$ |
| **Mean $COP_y$ Displacement** | 'Closed-Eyes/Compliant surface' | 0.381 | $p > 0.1$ |
| **Maximum $F_x$** | 'Open-Eyes/Compliant surface' | 0.403 | $p > 0.3$ |

### Classifier performance

SVM, MLP, K-NN and NB classifiers were employed to evaluate the performance of the 18 selected features. The selected features were kept in their discretized form during this evaluation. Table 3 below illustrates the mean, SD and range in values from 25 runs of the classification model. The MLP performed best across all performacne metrics. The percentage of subjects correctly classified by the MLP was 80±7 followed by NB (77±8.5), SVM (75±7.6) and finally K-NN at 75±6.3. The ROC-AUC which indicates the model’s capacity to distinguish between risk groups was 77±8 for the MLP, 73±9.8 for NB, 71±8.2 for the SVM and 70±8 for K-NN. The precision of all models in classifying the high-risk group was quite high but varied widely from 61-96% in the case of MLP for example. This variability was also seen in the models recall of the high-risk group which was poor on average with 52±13.1. This resulted in the trade-off between sensitivity and specificity captured by the F1 score presenting with a high variability also. Performance metrics for the low-risk group were higher on average for all scores except precision but all experienced less variability than in the classification of the high-risk group.

**Table 3.**
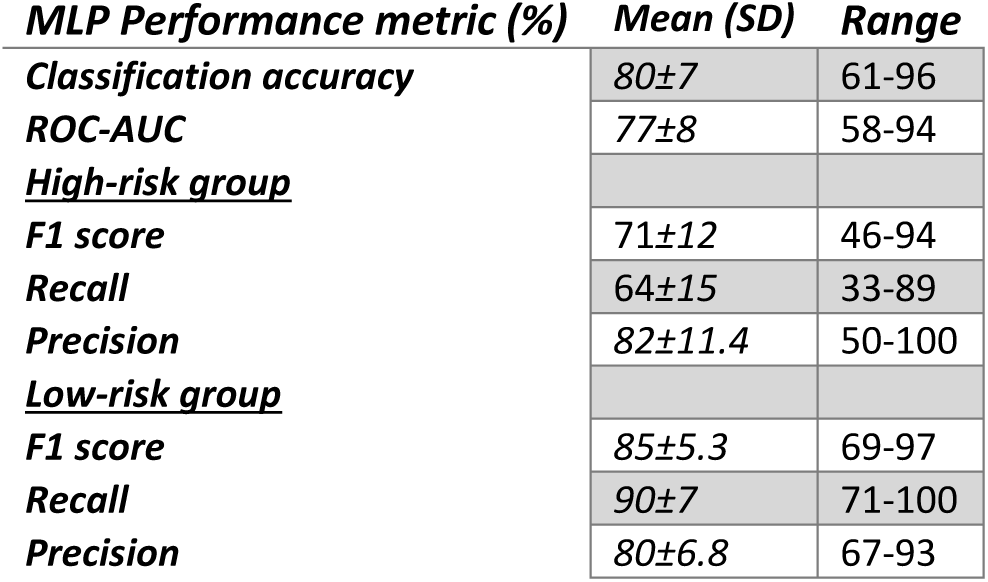

**Table 4.**
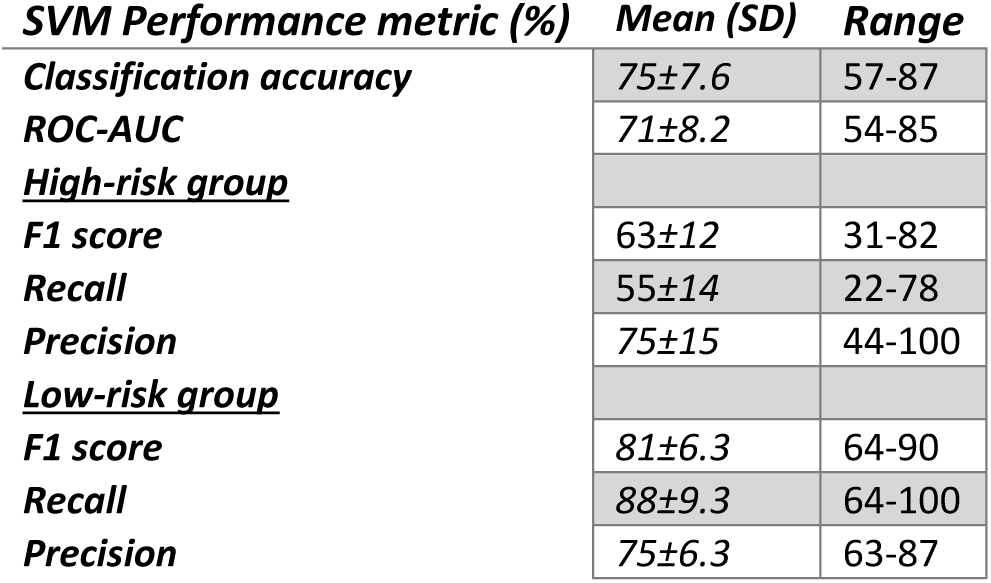

**Table 5.**

| **K-NN Performance metric (%)** | **Mean (SD)** | **Range** |
| --- | --- | --- |
| **Classification accuracy** | 75±6.3 | 67-93 |
| **ROC-AUC** | 70±8 | 58-92 |
| **<u>High-risk group</u>** |  |  |
| **F1 score** | 57±13 | 29-91 |
| **Recall** | 44±15 | 17-83 |
| **Precision** | 91±13.3 | 67-100 |
| **<u>Low-risk group</u>** |  |  |
| **F1 score** | 83±4.4 | 76-95 |
| **Recall** | 96±5.2 | 89-100 |
| **Precision** | 72.5±6 | 64-90 |

**Table 6.**

| **NB Performance metric (%)** | **Mean (SD)** | **Range** |
| --- | --- | --- |
| **Classification accuracy** | 77±8.5 | 67-93 |
| **ROC-AUC** | 73±9.8 | 58-92 |
| **<u>High-risk group</u>** |  |  |
| **F1 score** | 64±16 | 29-91 |
| **Recall** | 55±19 | 17-83 |
| **Precision** | 85±16 | 60-100 |
| **<u>Low-risk group</u>** |  |  |
| **F1 score** | 83±6.3 | 69-97 |
| **Recall** | 92±9.3 | 67-100 |
| **Precision** | 76±7.9 | 64-90 |

## Discussion

From the outset, variance in fall-risk predictability by force-plate measures of postural stability was recognized through conflicting results presented in numerous studies (21). Reasons proposed for this included the heterogeneity of the elderly population who may have various disorders and/or who may also have a stability deficit related to a specific component or combination of components of the postural control system (23,24). It was also stated how a significant number of static force-plate measures have been found to be capable of differentiating fallers vs. non-fallers, demonstrating the potential of some of these measures (17,18). Nonetheless studies combining stabilometry measures as part of a classification model have focused mostly on dynamic measures while thorough feature selection protocol have scarcely been investigated. This study aimed to use a thorough, evidence-based approach to feature selection which would lead to a more reliable classification accuracy of fall-risk in older subjects when using static force-plate measures. 18 features were selected based on a several-fold process of elimination which involved SVM-RFE, GA and ReliefF methods and finally analysed based on complementarity to ensure cohesion between features using SAFE.

The final classification models on average performed moderately well with the MLP seeing the highest average classification accuracy of 80% and an average ROC-AUC of 77% while K-nearest neighbours was the least accurate on average at 75% for along with a ROC-AUC of 70%. However, all final classification models were lacking in reliability with high variability in performance experienced. This variance was mainly centred around the high-risk group with poor recall scores across all models. Recall measures the sensitivity of the model, an important parameter in this case as a high rate of true-positive classifications are necessary for early fall-risk identification. A high precision was found for this group but with a large variation also, possibly indicating the classifications model’s capability to guess to a high accuracy. Approximately 40% of the sample population were allocated to the high-risk group which is representative of the elderly population (8). The number of medications, disorders, TMT scores and BesTest scores were not significantly different across groups indicating an equal contribution in variability from these sources from both groups. Imbalanced datasets can cause sub-optimal classifier performance however the extent of the imbalance within this dataset was mild and parameters were set during feature selection and in classifiers to compensate for this slight imbalance (44).

This issue of high bias/variance is common in machine-learning and is captured in what statistical-learning literature refers to as the ‘*bias-variance trade-off*’ (45,46). This trade-off can be represented on a continuum where on one end the model makes over-simplified assumptions (bias) about the training data and thus may not capture the underlying relationships sufficiently whereas on the other end the model is overly flexible and over-fitted to the datapoints in the training set, making generalization highly variable. High variability was seen across all classification indicating that high classification accuracies could not be achieved without under/over-fitting present. Optimization of the ‘*bias-variance trade-off*’ is mediated by the complexity of the concept being learnt, fall-risk in older subjects (45). It is likely that despite a thorough feature selection process, the limited information inherent in the static force-plate measures could not be compensated for and thus the model was insensitive to crucial components of fall-risk. This reveals a key limitation in static force-plate measures and may explain the conflicting conclusions made in previous studies on these measures’ predictability of fall-risk (21). The inclusion of dynamic measures of stability in such a protocol may improve sensitivity and overall reliability of classifications.

In the face of unreliable classifications, it is important to note that the feature selection methods used in this study, as in other similar studies, have demonstrated an impressive ability to identify useful features relevant to the target concept (32,38,47). The implementation of several methodologies also clearly benefited the final feature subset, exemplifying the benefits of a combinational approach. SVM-RFE removed most features from the original dataset, making feature selection more manageable for the GA and reducing the effect of redundancy further downstream. The optimal feature set included various notable measures that have been recognized in the research as relevant to balance performance e.g. F_Z_(SD), FD-CC and Mean COP_Y_ displacement while other measures warranting more investigation were also identified such as Maximum T_Z_and Freq95%-F_Y_ (21,48,49). A GA selects the most relevant features when all interactions in the subset are considered (50). Features identified by the GA in this study included all the features with a significant difference between risk-groups in the final subset. It is reasonable to regard many of the features chosen by GA as the most differential manifestations of postural instability in this sample. ReliefF identified features specifically sensitive to COP migration during the ‘*Open-Eyes/Firm surface*’ only. This method takes a different approach to feature selection to that of GA as it determines feature relevancy primarily based on interactions with other features while discriminative ability between classes is considered (3,51). Only two features were identified in both methods with ReliefF appearing to capture more intrinsic deteriorations in postural stability, i.e. Open/Closed-loop postural control during the condition where interactions between measures would be least perturbed. This is encouraging as the application of such methods to dynamic measures of stability may prove beneficial in revealing more relevant features for fall-risk classification.

## Limitations

Discrepancies between falls efficacy and the true balance performance capacity of an individual have been reported and may have affected the accuracy of fall-risk group allocation as subjects may not have displayed physical signs of balance impairment at the time of assessment but were identified as at risk. This discrepancy has however been shown to result in actual higher rates of fall-risk prospectively (52). The reasons for subject fall events were not recorded and as such it is possible that some falls may have been understandable given the conditions i.e. hazardous environment. The number of disorders and medications may not have been significantly different across groups however the severity and management of these factors may have been significantly different owing towards increased variability in the high-risk group.

## Conclusion

A thorough, evidence-based feature selection procedure was insufficient in improving the reliability of fall-risk classification and preventing under/over-fitting when using static force-plate measures only. The variability in classification performance was attributed to poor sensitivity in high-risk subjects, a crucial component in the assessment of early fall-risk detection revealing a key limitation in the measures used. The prescribed feature selection methods showed promise in identifying various measures relevant to fall-risk making the inclusion of dynamic measures in such a protocol potentially beneficial.

## Abbreviations

COP: Centre-of-pressure
COP_X_: Centre-of-pressure in Anterior-posterior direction
COP_Y_: Centre-of-pressure in Medio-lateral direction
F_X_: Anterior-posterior ground reaction force
F_Y_: Medio-Lateral ground reaction force
F_Z_: Vertical ground reaction force
GRF_s_: Ground reaction forces
FES: Falls efficacy scale (Short international version)
RMS: Root-Mean Square
SVM-RFE: Support vector machines-Recursive feature elimination
SVM: Support vector machines
NB: Naïve Bayes classifier
K-NN: K-Nearest neighbors’ classifier
SAFE: Self-adapting feature evaluation
GA: Genetic algorithm
MLP: Multi-layer perceptron

